# Defective synaptic plasticity in a model of Coffin-Lowry Syndrome is rescued by simultaneously targeting PKA and MAPK pathways

**DOI:** 10.1101/2022.06.07.495143

**Authors:** Rong-Yu Liu, Yili Zhang, Paul Smolen, Leonard J. Cleary, John H. Byrne

**Affiliations:** Department of Neurobiology and Anatomy, W.M. Keck Center for the Neurobiology of Learning and Memory, McGovern Medical School at the University of Texas Health Science Center at Houston, 6431 Fannin Street, Suite MSB 7.046, Houston, TX 77030

## Abstract

Empirical and computational methods were combined to examine whether individual or dual-drug treatments can restore the deficit in long-term synaptic facilitation (LTF) of the *Aplysia* sensorimotor synapse observed in a molecular model of Coffin-Lowry Syndrome (CLS). The model was produced by pharmacological inhibition of p90 ribosomal S6 kinase (RSK) activity. Simultaneous treatment with an activator of the mitogen-activated protein kinase (MAPK) isoform ERK and an activator of protein kinase A (PKA) resulted in enhanced phosphorylation of RSK and LTF to a greater extent than either drug alone and greater than their additive effects. Indeed, the combined drugs exerted synergistic effects on both RSK activation and LTF, fully restoring RSK phosphorylation and LTF. The extent of synergism appeared to depend on another MAPK isoform, p38 MAPK. Inhibition of p38 MAPK facilitated serotonin (5-HT)-induced RSK phosphorylation, indicating that p38 MAPK inhibits activation of RSK. Inhibition of p38 MAPK combined with activation of PKA synergistically activated RSK.

## INTRODUCTION

Coffin-Lowry Syndrome (CLS) is a genetic disorder caused by X-linked mutations in *rsk2*, and is characterized by cognitive impairment and craniofacial and skeletal abnormalities ^1^. In mammals, gene deletion studies have shown an essential role for *rsk2* in cognitive functions and learning ^2^. In several animal models, *rsk2* expresses the p90 isoform of RSK, a downstream effector of the ERK isoform of MAPK, phosphorylating and activating transcription activators such as CREB ^3-7^. Previous research has shown that memory formation in vertebrates and invertebrates requires CREB activation. For example, an *rsk2* null mouse model is deficient in several forms of long-term memory, and in stimulus-induced CREB phosphorylation ^2,8,9^. In *Aplysia*, inhibition of RSK activity by the membrane permeable molecule BI-D1870 (BID) significantly reduced serotonin (5-HT)-mediated CREB1 phosphorylation, impairing LTF and long-term enhancement of excitability (LTEE) ^10^, two independent cellular mechanisms for memory storage ^11^. These data suggest RSK is required for both LTF and LTEE and acts, at least in part, via activation of CREB1. LTF in *Aplysia* neurons may thus constitute a useful cellular model to study mechanisms underlying synaptic plasticity deficits in CLS ^10^. PKA and ERK are also essential for LTF ^11-13^ and contribute to the activation of RSK ^10,14^. These findings suggest that pharmacological activation of these kinases may enhance LTF and long-term memory (LTM). Activation in combination may synergistically (supra-additively) enhance LTF and LTM, because of complex interactions of kinase cascades critical for LTF and LTM, including feed forward and feedback loops. Such complexity contributes to nonlinearity of cellular responses to stimuli. two drugs targeting distinct kinase cascades could activate these auxiliary pathways and produce a nonlinear response to the stimulus, resulting in synergism. Rolipram, an inhibitor of cAMP phosphodiesterase (PDE), activates PKA and improves long-term potentiation (LTP) and LTM in normal rodents and in rodent models of Rubinstein-Taybi syndrome and traumatic brain injury ^15,16^. When rolipram is combined with the ERK pathway activator NSC295642 (NSC), weak LTF induced by 1 μM 5-HT is modestly enhanced, indicating the potential advantages of combining manipulations that target the PKA and ERK pathways ^17^. In order to apply this finding to CLS, we used 50 μM 5-HT to produce greater LTF that could then be reduced by RSK inhibition. We demonstrated that combined administration of rolipram and NSC was sufficient to restore LTF, and the two drugs acted synergistically.

## MATERIALS AND METHODS

### Cell culture and pharmacological treatment

Isolated SNs or SN-MN co-cultures from *Aplysia californica* (NIH *Aplysia* resource facility, University of Miami, Miami, FL) were prepared according to conventional procedures ^10,18^. The Standard 5-HT protocol is to treat SN cultures or SN-MN co-cultures with five, 5-min pulses of 50 μM 5-HT (Sigma, H9523) with a uniform interstimulus interval (ISI) of 20 min (onset to onset) ^18^. A separate group of control cultures were not treated with 5-HT, but were treated with vehicle alone (Veh) (L15:ASW) with ISIs of the Standard protocol.

BI-D1870 (BID, Santa Cruz, CAS 501437-28-1) is a small cell-permeant molecule that specifically inhibits RSKs ^10^. To activate the ERK pathway, we used the DUSP6 inhibitor NSC 295642 (Santa Cruz, CAS 77111-29-6). To activate the PKA pathway, we used rolipram (Sigma, R6520), an inhibitor of cAMP phosphodiesterase (PDE). SB 203580 was used to block p38 MAPK activity (Sigma, #559389). For drug treatment, SN or co-cultures were exposed to drugs for 30 min before and throughout 5-HT treatment. See supplemental materials for details.

### Immunofluorescence

SNs were starved with a solution of 50% L15 and 50% artificial seawater for 2 hours before drug treatment. After 5, 5-HT treatment, SNs were fixed for immunofluorescence at 1 h after 5-HT treatments following standard procedures ^10,19,20^. The antibodies against phosphorylated p90 RSK [anti-pRSK (Thr573), Cell Signaling, Cat # 9346] was used in 1:400 dilution. These experiments were performed in a blind manner so that the investigator analyzing the images was unaware of the treatment the SNs received. The number of samples (n) reported in Results indicates numbers of dishes assessed.

### Electrophysiology

Excitatory postsynaptic potentials (EPSPs) were recorded from MNs from the SN-MN co-cultures following established procedures^18,20-23^. The number of samples (n) reported in Results indicates the number of co-cultures. All the above experiments were performed in a blind manner so that the investigator performing the electrophysiology was unaware of the treatment the neurons received.

### Statistical analyses

#### For immunocytochemistry

SNs were isolated from the same animal in each experimental repetition (Figs. 4, 6, and 7). When SNs were fixed for immunocytochemistry followed by confocal imaging, the imaging parameters were fixed throughout each experiment repetition. In this case, because the data has these known relationships, repeated measures is the appropriate statistical analysis. Therefore, repeated measures one-way (RM) ANOVA was used on raw data, followed by the post hoc Student-Newman–Keuls method (SNK) for multiple comparisons analysis ^10,17,20-22,24,25^. SigmaPlot version 11 (Systat Software, Inc.) was used to perform all statistical analyses.

#### For electrophysiological experiments

the amplitudes of the EPSPs were assessed before (pre-test) and 24 h after 5-HT treatment (post-test). In contrast to immunofluorescence, these data were normalized for one-way ANOVA analysis, followed by SNK post hoc analysis. Post-test data were normalized to the corresponding measurements made at pre-test. To examine possible changes in resting potentials and input resistance of MNs and SNs caused by BID, measurements were made before (pre-test) and 24 h after vehicle or BID treatment (post-test). Student’s t-tests was used to compare the changes [(post-pre)/pre)] in the BID group with those in the vehicle group.

To evaluate synergistic effects of dual drug treatments on LTF and RSK activation, we used a two-way ANOVA to assess the interaction between two treatment factors ^26-29^. The significance of the combined-drug interaction was used to evaluate the null hypothesis that the combined effect of two drugs is merely additive (Fig. 3, Figs. 4B3 and C3, and Fig. 6C3).

Data from all experiments were presented as means ± SEM, and p < 0.05 was considered to represent statistical significance.

## Results

### 1. NSC and rolipram alone failed to restore impaired LTF

We first examined whether LTF impaired by BID can be rescued by NSC or rolipram alone. Normal LTF was induced by exposing *Aplysia* sensory neuron – motor neuron (SN-MN) co-cultures to 5 pulses of 5-HT at standard concentration (50 μM, 20 min interstimulus interval (ISIs)) ^30^. EPSPs were measured prior to 5-HT treatment (pre-test), and 24 h after treatment (post-test) in six groups: 1) 5-HT, 2) BID+5-HT, 3) Rolipram (R)+5-HT (R+5-HT), 4) R+BID+5-HT, 5) NSC+5-HT, 6) NSC+BID+5-HT (Fig.1). The standard protocol led to a 51 ± 6% (n = 10) increase in EPSP amplitude, whereas BID reduced LTF to 12 ± 8% (n = 9). 0.1 μM Rolipram or 0.01 μM NSC alone each produced increases in LTF. The EPSP increase in the R+5-HT group was 62 ± 9% (n = 10) and 64 ± 6% in the NSC+5-HT group (n = 10). Similarly, these two drugs alone slightly enhanced BID-impaired LTF. The EPSP increase in the R+BID+5-HT group was 35 ± 10% (n = 9) and 29 ± 6% in the NSC+BID+5-HT group (n = 10). A one-way ANOVA indicated significant overall differences among the groups (F_5,52_ = 7.08, p < 0.001). However, post hoc comparisons revealed that neither rolipram alone nor NSC alone significantly enhanced LTF (5-HT vs. R+5-HT, q = 1.502, p = 0.293; 5-HT vs. NSC+5-HT, q = 1.825, p = 0.407). Although BID significantly reduced LTF (5-HT vs. BID+5-HT, q = 5.001, p < 0.05), neither rolipram nor NSC significantly enhanced BID-impaired LTF (BID+5-HT vs. R+BID+5-HT, q = 2.892, p = 0.112; BID+5-HT vs. NSC+BID+5-HT, q = 2.152, p = 0.134).

**Figure 1.**
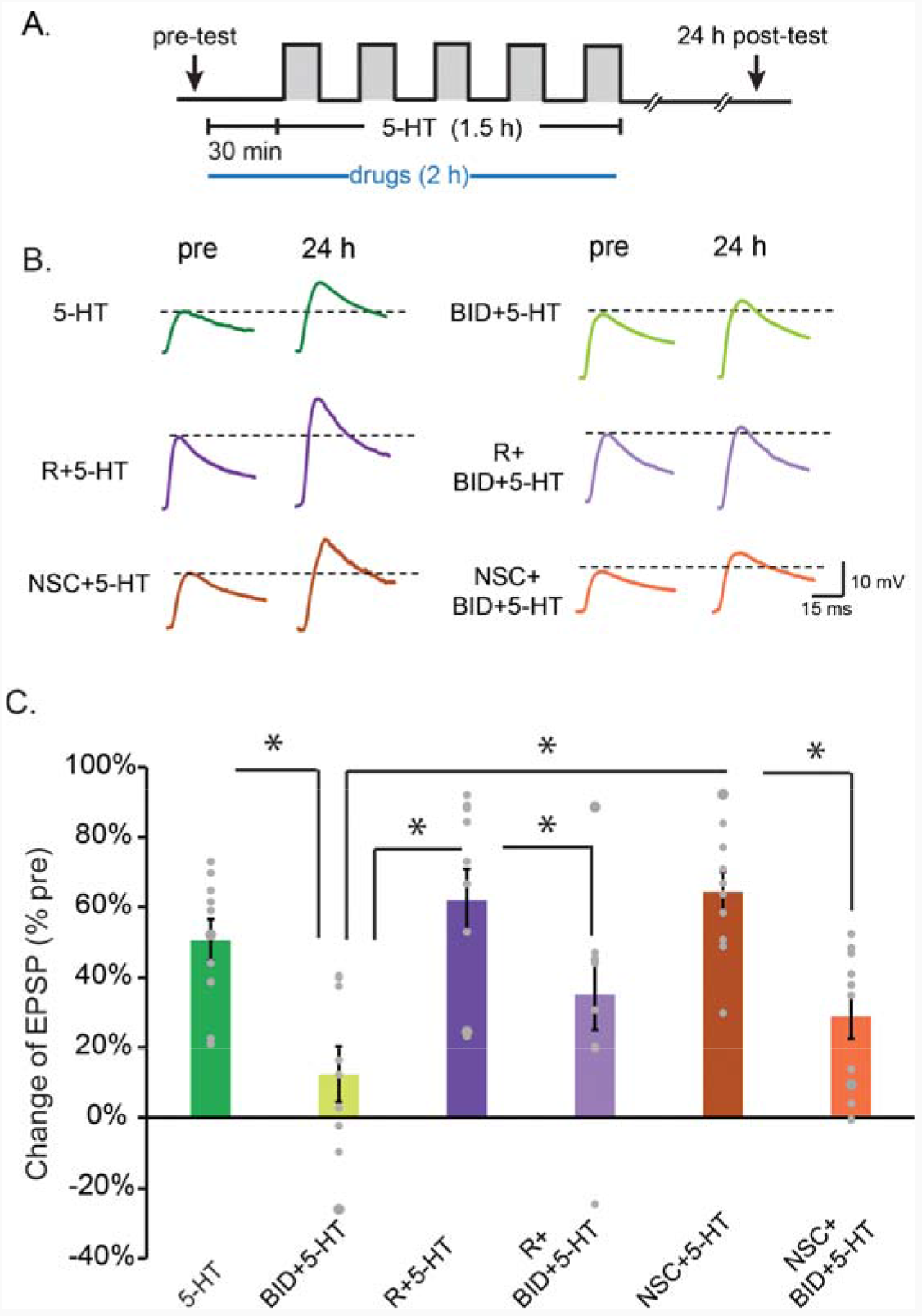
Individual effects of NSC and rolipram on impaired LTF induced by pharmacological inhibition of RSK. **A**, Protocol for standard 5 pulses of 5-HT (50 μM) treatment, in the presence of BID, with NSC and/or rolipram (R) application. **B**, Representative EPSPs recorded from MNs in SN-MN co-cultures before (Pre) and 24 h after 5-HT treatment. **C**, Summary data. Inhibition of RSK activity impaired 5-HT-induced LTF. This impairment was not rescued by NSC or R alone. In this and subsequent illustrations, bar height represents the mean, small bars represent standard error of the mean (SEM), and significant differences are indicated by * for p < 0.05.

### 2. Combining NSC and rolipram rescued LTF

SN-MN co-cultures were pre-incubated with BID in the presence of NSC and/or rolipram for 30 min, then treated with five pulses of 5-HT in continued drug presence (Fig. 1A). Four groups of SN-MN co-cultures were assessed: 1) 5-HT group to assess normal LTF, 2) BID+5-HT group, 3) NSC+R+5-HT group to assess normal LTF enhanced by combined drugs, and 4) NSC+R+BID+5-HT group to assess rescue of LTF by combined drugs (Fig. 2). Standard 5-HT treatment led to a 49 ± 6% (n = 13) increase in EPSP amplitude, whereas BID reduced LTF to 15 ± 6% (n = 11). Combining NSC with rolipram increased normal LTF to 94 ± 13% (n = 9). In the NSC+R+BID+5-HT group, LTF was enhanced to 100 ± 18% (n = 10). One-way ANOVA indicated significant overall differences among the groups (F_3,39_ = 12.69, p < 0.001). LTF in the BID + 5-HT group was significantly less than that in the 5-HT group (q = 3.231, p = 0.028). Combined drugs significantly enhanced LTF (NSC+R+5-HT vs. 5-HT, q = 4.056, p = 0.007). Importantly, LTF in the NSC+R+BID+5-HT group was significantly greater than in the 5-HT group (q = 4.72, p = 0.005) and BID+5-HT group (q = 7.573, p < 0.001). Moreover, no significant difference in LTF was observed between the NSC+R+5-HT and NSC+R+BID+5-HT groups (q = 0.493, p = 0.729). Therefore, comparing Fig. 2 with Fig. 1, only the combined drugs fully restored LTF.

**Figure 2.**
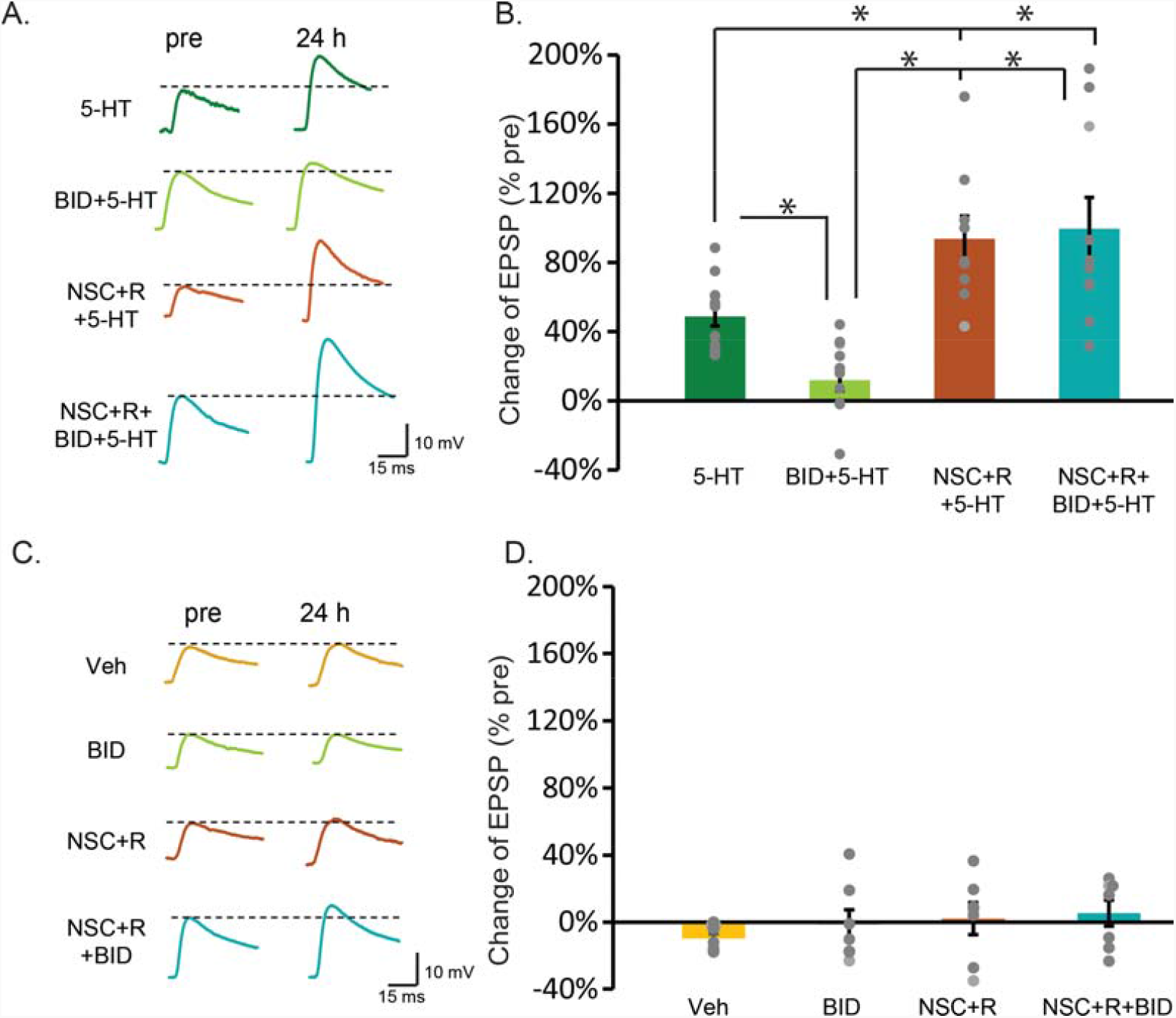
Effects of NSC and rolipram on impaired LTF and basal EPSPs. **A**, Representative EPSPs recorded from MNs in SN-MN co-cultures before (Pre) and 24 h after 5-HT treatment. **B**, Summary data. Inhibition of RSK impaired 5-HT-induced LTF. This impairment was fully rescued by co-applying NSC and R. **C**, Representative EPSPs in absence of 5-HT. **D**, Summary data indicate that BID, NSC, and R alone, or in combination, did not change basal synaptic strength.

Previous results suggested that these drugs could affect basal synaptic strength (Liu et al., 2017). Therefore, control experiments were performed to examine the extent to which potentiation of LTF could result from EPSP increases due to the drugs without 5-HT ^17^. SN-MN co-cultures were incubated with 0.01 μM NSC and 0.1 μM rolipram, with or without BID for 2 h. EPSP amplitudes with Veh alone did not change significantly at 24 h (−10 ± 3%, n = 7) (Fig. 2B). BID alone also did not change synaptic strength significantly (−1 ± 9%, n = 7). For NSC+R, EPSPs increased by 2 ± 10% (n = 7) at 24 h. In NSC+R+BID, EPSPs increased by 5 ± 8% (n = 7). A one-way ANOVA indicated no significant overall differences among the groups (F_3,24_ = 0.721, p = 0.549). Thus, NSC and rolipram did not affect basal synaptic strength. Therefore, the observed enhancement in LTF (Fig. 2A) was due to the interaction between the PKA and ERK pathways activated by drugs and 5-HT.

### 3. Analysis of synergism for the effect of the drug combination on LTF

The above findings that the combined, but not individual, drugs fully rescued BID-impaired LTF supports the hypothesis that there are supra-additive or synergistic effects between ERK activation by NSC and PKA activation by rolipram. The average normal LTF was 51 ± 6% in Fig. 1 and 49 ± 6% in Fig. 2, and BID-impaired LTF was 12 ± 8% in Fig. 1 and 15 ± 6% in Fig. 2. Two-sample t tests indicated the changes in EPSP were not significantly different between the normal (t_1,21_ = 0.059, p = 0.81), or BID-impaired (t_1,18_ = 0.061, p = 0.81) groups. Therefore the responses to 5-HT or BID +5-HT in Figs. 1 and 2 were combined, into a normal LTF group [Fig. 3A, NSC(−)R(−)] and a BID-impaired LTF group [Fig. 3B, NSC(−)R(−)], to compare individual drug effects to combination effects. The analysis of normal LTF included four groups: 1) 5-HT; 2) R+5-HT, 3) NSC+5-HT, and 4) NSC+R+5-HT. Normal LTF was 50 ± 4%, n = 23 (Fig. 3A, triangle symbol). Rolipram produced a small additional increase to 62 ± 9% (n = 10) (Fig. 3A, square). NSC also caused a small increase to 64 ± 6% (n = 10) (Fig. 3A, asterisk). Combining NSC and rolipram increased LTF to 94 ± 13% (n = 9) (Fig. 3A, oval). To analyze whether NSC and rolipram acted synergistically, we performed a two-way ANOVA as suggested by Slinker (1998) and applied by multiple groups ^27-29^. Two drugs are deemed to act synergistically if the F statistic for the interaction effect is less than the critical value of 0.05. If the interaction between the two drugs fails to reach significance, the effects are deemed non-synergistic. A two-way ANOVA performed with the data of Fig. 3A showed a significant main effect of NSC (F_1,48_ = 5.061, p = 0.029), and a trend of enhancement, but not statistically significant, effect of rolipram (F_1,48_ = 3.954, p = 0.052). More important, no significant interaction between rolipram and NSC (F_1,48_ = 1.423, p = 0.239) was observed, indicating an additive but not a synergistic interaction between these drugs in their effects on normal LTF.

**Figure 3.**
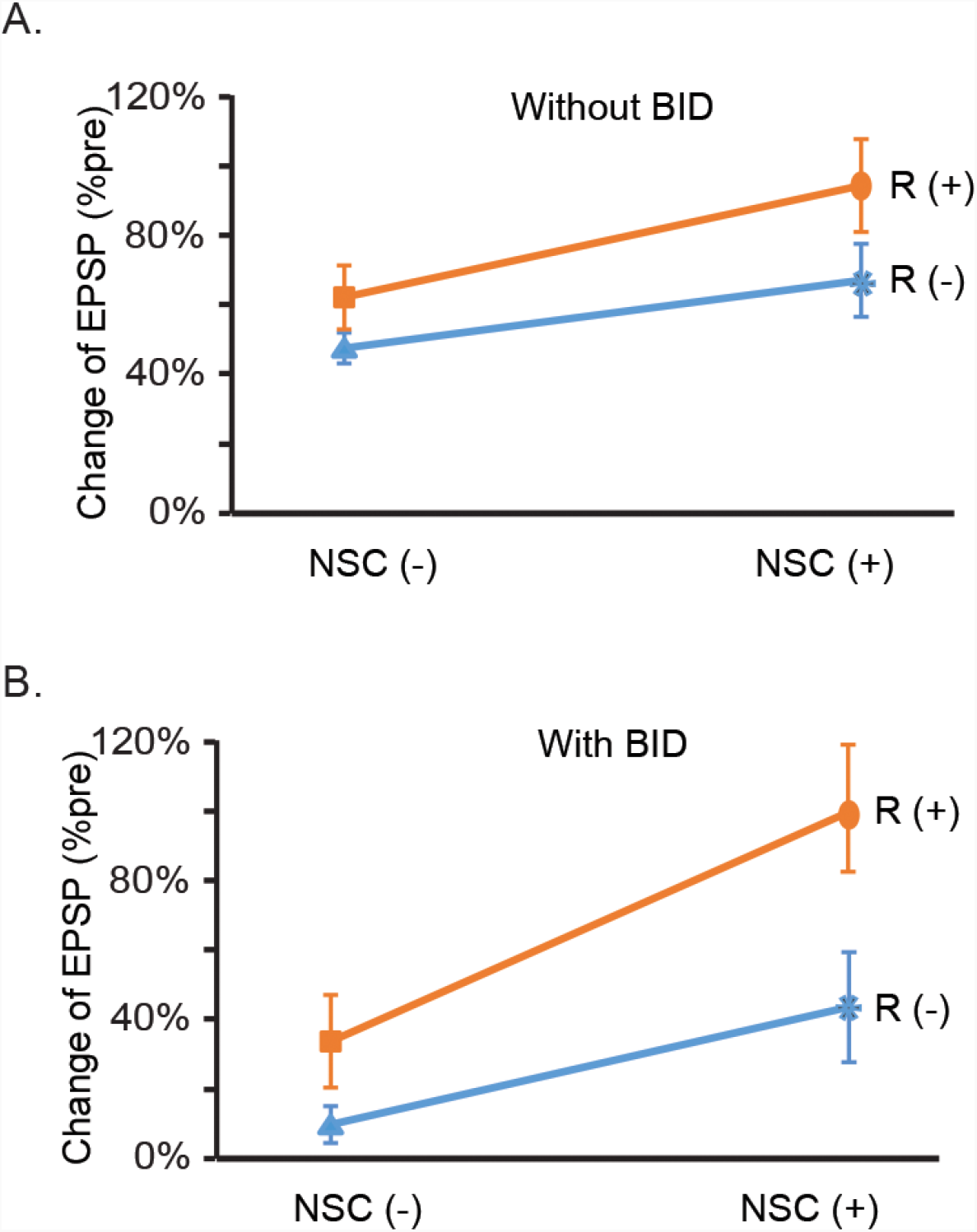
Tests of synergism for the drug combination using the data from Figs. 1 and 2. **A**, Drug effects of NSC and R on LTF in the absence of BID. Statistical analysis using two-way ANOVA revealed no significant interaction between the groups and therefore a lack of synergism between NSC and rolipram on normal LTF. **B**, Drug effects on BID-impaired LTF. Statistical analysis using two-way ANOVA revealed a significant interaction between rolipram and NSC, indicating synergistic rescue of impaired LTF.

The analysis of the impaired LTF in the CLS model included: 1) BID+5-HT, 2) NSC+BID+5-HT, 3) R+BID+5-HT, and 4) NSC+R+BID+5-HT (Fig. 3B). The impaired LTF in BID+5-HT was 14 ± 5% (n = 20) (Fig. 3B, triangle symbol). The application of rolipram produced increase in EPSP to 35 ± 10% (n = 9) (Fig. 3B, square), and NSC increased the EPSP by 29 ± 6% in the NSC+BID+5-HT group (n = 10). Combining NSC and rolipram with 5-HT increased LTF to 99 ± 18% (n = 10) (Fig. 3B, oval). A two-way ANOVA revealed that the enhancements from rolipram or NSC alone were both significant (Factor R, F_1,45_ = 21.49, p < 0.001; Factor NSC, F_1,45_ = 16.05, p < 0.001). Importantly, the interaction between these two drugs was also significant (F_1,45_ = 6.159, p = 0.017), confirming that addition of the two drugs together has a synergistic effect to rescue impaired LTF in the cellular CLS model.

### 4. Combined drugs synergistically activated RSK

We investigated molecular mechanisms that may explain, at least in part, this combined-drug synergism. As discussed above, PKA and ERK both activate RSK ^14^. Therefore, the synergistic effect of the drugs on LTF may be due to synergistic enhancement of RSK phosphorylation. To test this hypothesis, we measured levels of phosphorylated RSK (pRSK) with immunofluorescence. Rolipram and/or NSC was applied, with 5-HT, to isolated SNs and RSK phosphorylation was assessed 1 h after 5-HT treatment (Fig. 4A). pRSK levels in 5-HT, R+5-HT, NSC+5-HT and R+NSC+5-HT groups were normalized to the group treated with vehicle control. Compared to Veh, 5-HT led to a 23 ± 7 % increase in pRSK. The increase of pRSK levels in the R+5-HT and NSC+5-HT groups was 21 ± 4% and 18 ± 4%, respectively. However, the increase of pRSK in NSC+R+5-HT was 55 ± 10 % (Figs. 4B1-B3). One-way RM ANOVA indicated significant overall differences among the groups (F_3,15_= 8.42, p = 0.002). Post hoc analysis indicated the pRSK level in NSC+R+5-HT was significantly greater than in the other groups (NSC+R+5-HT vs. 5-HT, q = 5.06, p = 0.003; NSC+R+5-HT vs. R+5-HT, q = 5.82, p = 0.003; NSC+R+5-HT vs. NSC+5-HT, q = 6.44, p = 0.002). No significant difference in pRSK was observed between NSC 5-HT vs. 5-HT (q = 0.76, p = 0.855), or R+5-HT vs. 5-HT (q = 0.176, p = 0.903). To examine whether the dual drug interaction was synergistic, a two-way RM ANOVA (Methods) was performed on the 5-HT, R+5-HT, NSC+5-HT, and R+NSC+5-HT groups. This, however, revealed no significant interaction between rolipram and NSC (Fig. 4B3) (F_1,5_ = 1.428, p = 0.444).

**Figure 4.**
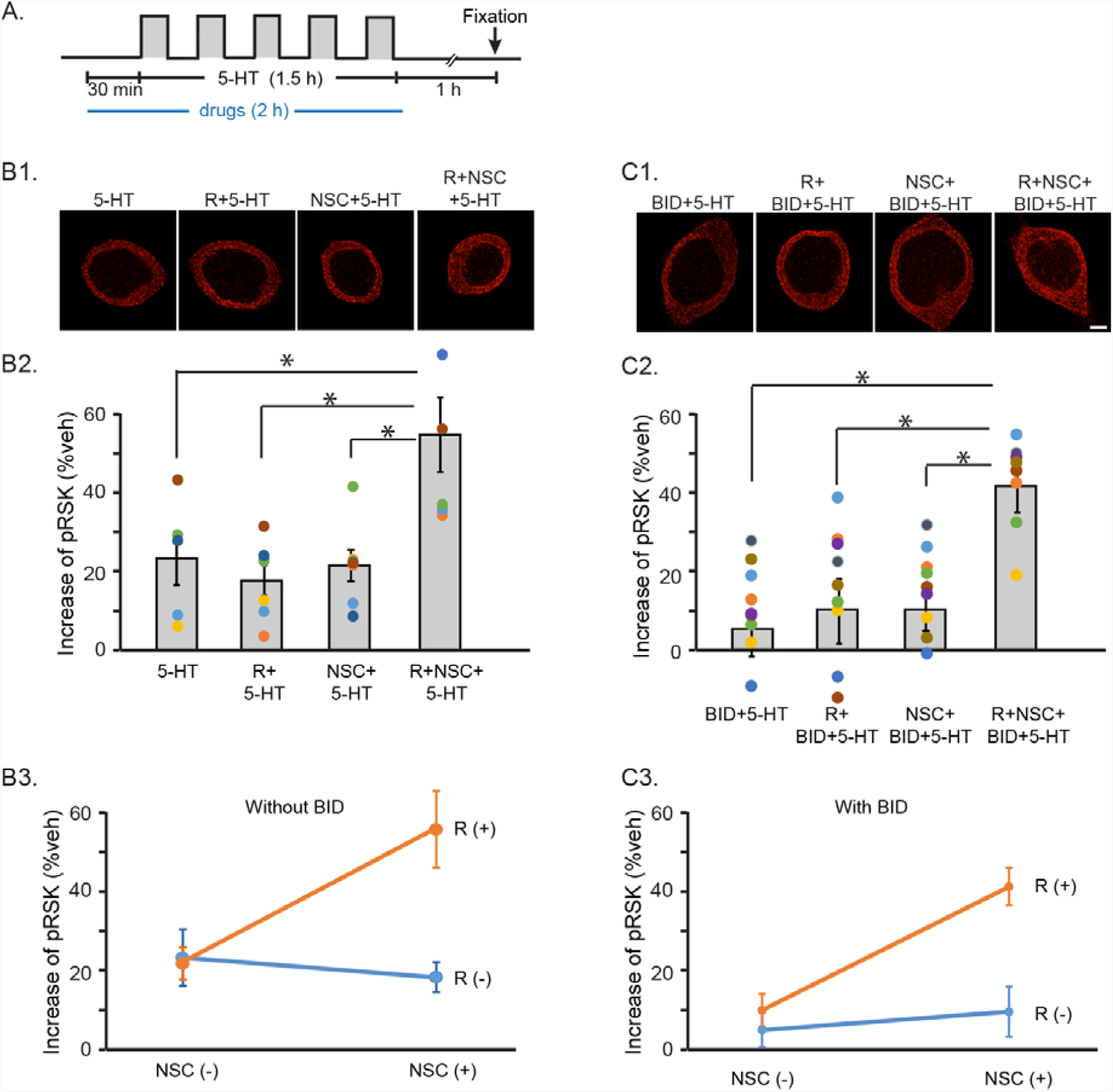
Effects on RSK activation of individual and combined drugs. **A**, Protocol for standard 5 pulses of 5-HT (50 μM) treatment, with NSC and/or R application, without (B) or with (C) BID. **B**, Without BID. **B1**, representative confocal images of pRSK in SNs 1 h after 5-HT treatment, with a single drug or combined drugs. Scale bar, 20 μm. **B2**, Summary data. Dual drugs potentiated the 5-HT-induced increase in pRSK. **B3**, Two-way ANOVA indicated no synergism between R and NSC. **C**, With BID. **C1**, representative confocal images of pRSK in SNs 1 h after 5-HT treatment, with a single drug or combined drugs. **C2**, Summary data. A single drug slightly restored pRSK, and dual drugs substantially increased 5-HT-induced, BID-impaired pRSK. **C3**, Two-way ANOVA revealed a synergistic interaction between NSC and R. Different colored dots in each group indicate individual experiments and the same color in different groups represent one experiment. Scale bar, 20 μm.

Next, we investigated possible synergism between rolipram and NSC in restoring 5-HT-induced pRSK impaired by BID. Rolipram and/or NSC was applied to isolated SNs in the presence of BID, and the effects on RSK phosphorylation were examined 1 h after 5-HT treatment (Figs. 4C1-C3). Immunoreactivities to pRSK in the BID+5-HT, R+BID+5-HT, NSC+BID+5-HT and R+NSC+BID+5-HT groups were normalized to vehicle control. Compared to Veh, in the presence of BID, 5-HT only led to a 5± 4 % increase in pRSK. The increase in pRSK in the R+BID+5-HT and NSC+BID+5-HT group was 11 ± 4% and 10 ± 6% of control, respectively. However, the increase of pRSK in R+NSC+BID+5-HT was 42 ± 5 % of control. One-way ANOVA indicated significant overall differences among the groups (F_3,31_= 10.67, p < 0.001). Post hoc comparison indicated pRSK in the NSC+R+BID+5-HT group was significantly greater than in the other groups (NSC+R+BID+5-HT vs. BID+5-HT, q = 7.20, p < 0.001; NSC+R+BID+5-HT vs. R+BID+5-HT, q = 6.20, p < 0.001; NSC+R+BID+5-HT vs. NSC+BID+5-HT, q = 6.28, p < 0.001). Moreover, no significant difference in pRSK was observed between NSC+BID+5-HT vs. BID+5-HT (q = 0.95, p = 0.506), or R+BID+5-HT vs. BID+5-HT groups (q = 1.032, p = 0.748).

To determine whether the dual drug interaction was synergistic, a two-way RM ANOVA was performed on the BID+5-HT, R+BID+5-HT, NSC+BID+5-HT and R+NSC+BID+5-HT groups. This revealed a significant (F_1,7_ = 7.488, p = 0.029) interaction between rolipram and NSC (Fig. 4C3), indicating a synergistic enhancement of the BID-impaired increase in pRSK.

### 5. Computational modeling of synergism

PKA activates RSK by multiple pathways, forming a positive feedforward loop (Fig. 5A, pathways 10, 3->6->7). It is plausible that some combinations of drug concentrations would take advantage of the nonlinear interactions between these pathways to act synergistically. Although synergism was observed empirically, it was only evident when RSK was partially inhibited (i.e., the CLS model). To gain insights into the molecular determinants of synergism we exploited a previously developed computational model that simulated the 5-HT-induced dynamics and interactions of kinase cascades and growth factors (Fig. 5A)^14^. Modeling can assess the effectiveness, and possible synergism, of a large number of combinations of NSC and rolipram (with or without an RSK inhibitor).

**Figure 5.**
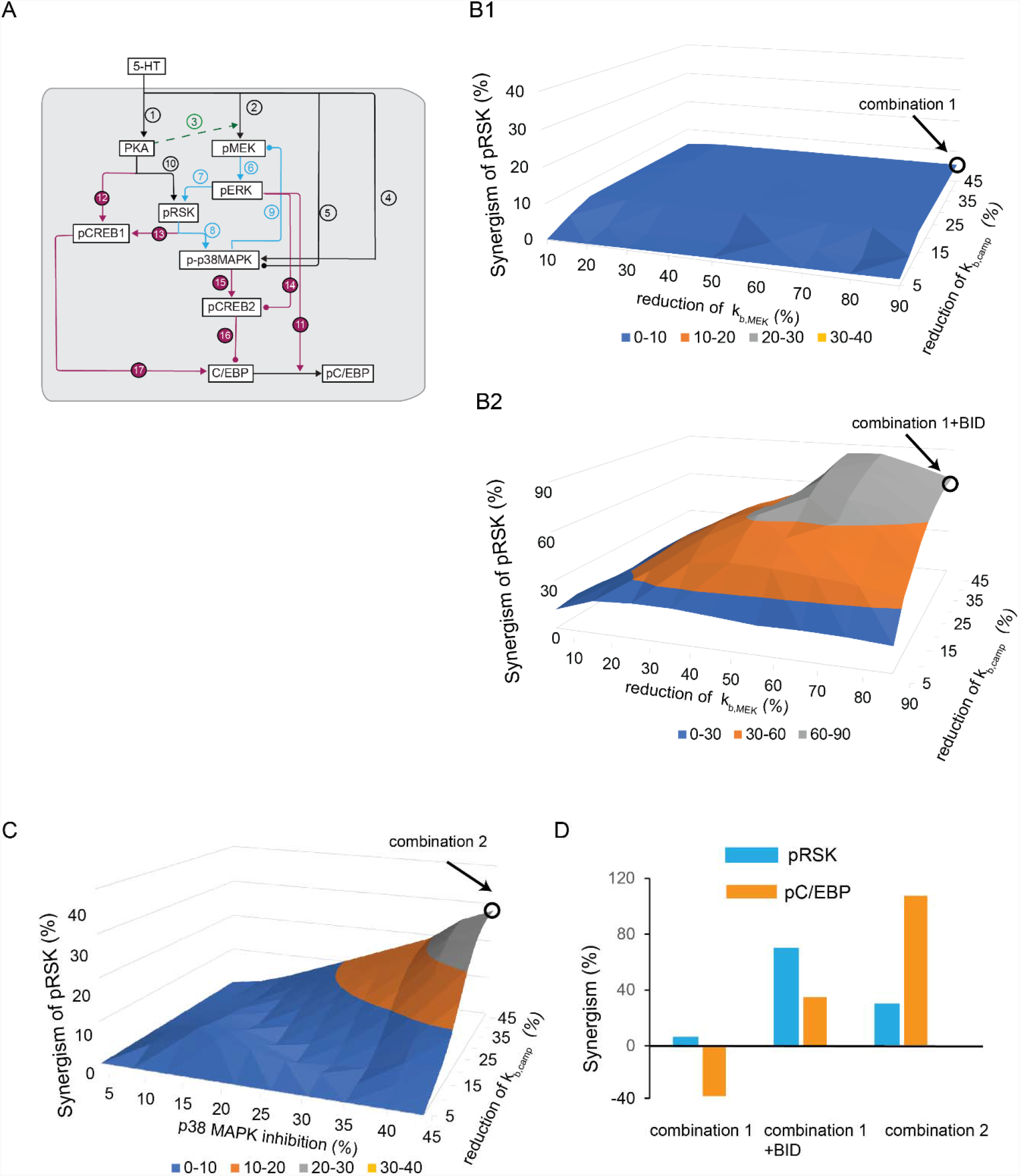
Computational modeling of synergism. **A**, Schematic model for molecular network of LTF, simplified from Zhang et al. (2021). The PKA and MAPK cascades interact to regulate the phosphorylation and activities of RSK. PKA, ERK, and RSK in turn regulates the phosphorylation and activation of CREB1 and CREB2. These effects on CREB1 and CREB2 induce the expression of immediate-early genes such as C/EBP. p38 MAPK is activated downstream of RSK, and in a negative feedback loop, p38 MAPK inhibits the activation of MEK, and therefore ERK and RSK. Arrowheads indicate activation, circular ends indicate repression. **B**, 3-D plot of synergism in pRSK induced by combined effects of cAMP phosphodiesterase inhibition (k_b,camp_ reduction) and ERK activation (*k*_*b,ME*K_ reduction), in the absence of BID (**B1**) or presence of BID (**B2**). **C**, 3-D plot of synergism in pRSK induced by combined p38 MAPK inhibition and cAMP phosphodiesterase inhibition. **D**, Comparison of synergism in pRSK (blue) and pC/EBP (orange) in different treatment combinations.

Model equations and parameters describing the PKA and MAPK pathways and their interactions remain as in Zhang et al. (2021) (see supplemental materials and Table S1, Eqs. 1-40). However, to simulate LTF, the synthesis and activation of the transcription factor CCAAT enhancer binding protein (C/EBP) was added (Fig. 5A, pathways 11, 16-17) (supplemental materials, Eqs. 41-43), given that C/EBP activation is essential for LTF ^31-34^. The expression of C/EBP is regulated by CREB1 and CREB2 (Fig. 5A, pathways 16-17), and C/EBP is activated by active, phosphorylated ERK (pERK) (Fig. 5A, pathway 11). The peak level of active, phospxhorylated C/EBP (pC/EBP) was used as a proxy for the strength of LTF. The effect of BID was represented by reducing the effects of RSK on CREB1 and p38 MAPK by 60% (*k*_*f,p38_RSK*_ in Eq. 24 and *k*_*RSK,CREB1*_ in Eq. 28, i.e., suppressing pathways 8 and 13 in Fig. 5A). The effect of NSC was represented by reducing the deactivation rate of MEK (*k*_*b,ME*K_ in Eqs. 3-4 and 35-36), the upstream kinase of ERK. The effect of rolipram was represented by reducing the degradation rate of the activator of PKA, cyclic AMP (cAMP) (*k*_*b,camp*_ in Eq. 11).

Because the model is deterministic, statistical approaches are not needed to assess simulated synergism. Consequently, synergism of the combined drugs was calculated as follows:

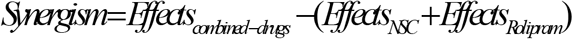

The variables ‘Effects’ represent the increases of pRSK or pC/EBP due to treatment by drugs and 5-HT, compared to 5-HT treatment only. *Effects*_*NSC*_ + *Effects*_*Rolipram*_ is the value of additive effect if NSC and rolipram act independently. We assumed that the combined drugs have synergistic effects as long as the value of the variable ‘Synergism’ is > 0 in the model, although a synergism of low value (e.g., <10%) might not be detectable in empirical studies. A larger score of Synergism represents stronger synergism.

NSC enhances pERK by inhibiting the dephosphorylation of MEK. However, the dose-response curve of NSC concentration vs. MEK dephosphorylation has not been empirically determined. Therefore, we simulated NSC effects by decreasing the dephosphorylation of MEK over a broad range. We decreased the dephosphorylation rate constant *k*_*b,ME*K_ by steps of 10%, from 100% down to 10% of its standard value. By reducing *k*_*b,ME*K_ to 10% of its standard value, pRSK was increased by ∼ 30%. For rolipram, we similarly decreased the degradation rate constant *k*_*b,camp*_ by steps of 5%, from 100% to 55% of its standard value. Compared to *k*_*b,ME*K_, a smaller step of change in *k*_*b,camp*_ was selected, because *k*_*b,camp*_ reduction was apparently more efficient in enhancing pRSK. By reducing *k*_*b,camp*_ to 55% of its standard value, pRSK was increased by ∼ 25%. The effect of each drug “pair” was simulated, for a total of 100 combinations (10 levels of *k*_*b,ME*K_ reduction combined with 10 levels of *k*_*b,camp*_ reduction). The X and Y axes in Figs. 5B-C denote the percentages of the standard values of these parameters. The Z axes show the Synergism variable. In the absence of BID, all combinations of treatments induced a pRSK Synergism score near 5%, thus the combined drugs can increase pRSK ∼5% more than the sum of individual drug effects (i.e., weak synergism, Fig. 5B1). We used the combination that produced the highest synergism for further simulations, which is a 90% *k*_*b,ME*K_ reduction combined with 45% *k*_*b,camp*_ reduction, denoted ‘combination 1’ (arrow in Fig. 5B1). In the absence of BID, combination 1 induced a Synergism score of -36% in the peak of pC/EBP, the proxy for the strength of LTF (Liu et al. 2013) (Fig. 5D). The negative score suggests the weak synergism in pRSK induced by ‘combination 1’ failed to induce synergism in pC/EBP.

The effect of BID was represented by reducing the effects of RSK on CREB1 and p38 MAPK by ∼50% (*k*_*f,p38_RSK*_ in Eq. 24 and *k*_*RSK,CREB1*_ in Eq. 28). Combination 1 then induced a strong Synergism score (69%) in pRSK (Fig. 5B2) and a Synergism score (34%) in pC/EBP (Fig. 5D). Thus, simulated BID treatment substantially increased the synergism in pRSK and pC/EBP.

The model also provided insights into the lack of rolipram and NSC synergism in enhancing normal LTF (i.e., Fig. 3A) and pERK elevation (Fig. 4B). Increased pRSK leads to increased p38 MAPK activity (Fig. 5A, pathway 8). We hypothesized that this lack of synergism by NSC and rolipram might be due, at least in part, to a suppressive effect of p38 MAPK on ERK. As shown in Fig. 5A (pathways 6 -> 7 -> 8 -> 9), and based on empirical findings from our laboratory (Zhang et al. 2017, 2021), the model includes a negative feedback loop in which RSK, activated by pERK, leads to p38 MAPK activation, which in turn inhibits activation of MEK and ERK, potentially limiting the extent of 5-HT-induced RSK activation and LTF. Moreover, enhanced p38 MAPK induced by combination 1 activates transcription inhibitor CREB2 (Fig. 5A, pathway 15), subsequently suppressing the expression of C/EBP (Fig. 5A, pathway 16). Thus, ‘combination 1’ leads to a negative Synergism score in pC/EBP (Fig. 5D). The addition of BID to suppress activation of p38 MAPK by pRSK (Fig. 5A, pathway 8) may suppress the negative feedback, substantially increasing synergism in pRSK. However, BID will also suppress the activation of CREB1 by pRSK (Fig. 5A, pathway 13), which would tend to limit the increase of synergism in the critical output variable pC/EBP (Fig. 5D).

Therefore, we predicted that an alternative drug combination to induce synergism might be rolipram and an inhibitor of p38 MAPK. A p38 MAPK inhibitor could relieve the inhibition of pRSK by p38 MAPK without affecting the activation of CREB1 by pRSK. To computationally test this hypothesis, we simulated the combined effects of p38 MAPK inhibition and rolipram on pRSK and pC/EBP. To simulate p38 MAPK inhibition we gradually decreased the downstream effects of p38 MAPK on MEK and CREB2 (*k*_*EP38, MEK*_ in Eq. 10 and *k*_*p38,CREB2*_ in Eq. 31, i.e., suppressing pathways 9 and 15 in Fig. 5A) by steps of 5%, from 100% down to 55% of the standard values. By reducing *k*_*EP38, MEK*_ to 55% of its standard value, pRSK was increased by ∼ 20%. At the same time, we gradually decreased the degradation rate constant of cAMP by steps of 5%, from 100% down to 55% of the standard values. Supporting the new hypothesis, simulations of these 100 pairs of parameter changes showed that combined p38 MAPK inhibition and k_b,camp_ reduction induced strong synergism in pRSK (Fig. 5C). We used the combination that produced the greatest synergism for further simulations, which was 45% p38 MAPK effect reduction combined with 45% k_b,camp_ reduction, denoted ‘combination 2’ (arrow in Fig. 5C). Combination 2 induced a higher Synergism score (28%) in pRSK, which in turn induced a strong Synergism score (106%) in pC/EBP (Fig. 5D). These results predict that an inhibitor of p38 MAPK in combination with rolipram will enhance normal LTF.

### 6. Inhibition of p38 MAPK potentiates phosphorylation of RSK

It is important to empirically test the model prediction (Fig. 5) that p38 MAPK activation can limit the extent of 5-HT induced RSK activation. Immunostaining for pRSK was performed 1 h after standard 5-HT treatment in the presence of the p38 MAPK inhibitor SB203580 (SB, Fig. 6B). The 5-HT-induced phosphorylation of RSK in SNs averaged a 15 ± 3% increase compared to vehicle control. SB itself produced little change in pRSK (−5 ± 4%). However, pRSK increased to 25 ± 4% of control in the SB+5-HT group. One-way RM ANOVA indicated significant overall differences among the treatment groups (F_3.21_ = 18.318, p < 0.001). Post hoc comparisons (SNK) revealed 5-HT induced significant increases in pRSK (5-HT vs. Veh, q = 4.402, P = 0.005; 5-HT vs. SB, q = 6.254, p < 0.001). Importantly, there was a significant difference between the SB+5-HT group and the other two groups (SB+5-HT vs. Veh, q = 7.614, p < 0.001; SB+5-HT vs. 5-HT, q = 3.212, *p* = 0.034; SB+5-HT vs. SB, q = 9.466, p < 0.001), indicating SB significantly enhanced activation of RSK. There was no basal effect from SB (no significant difference between Veh and SB, q= 1.851, p = 0.205). These results validate the prediction that p38 MAPK acts to limit the extent of RSK activation.

**Figure 6.**
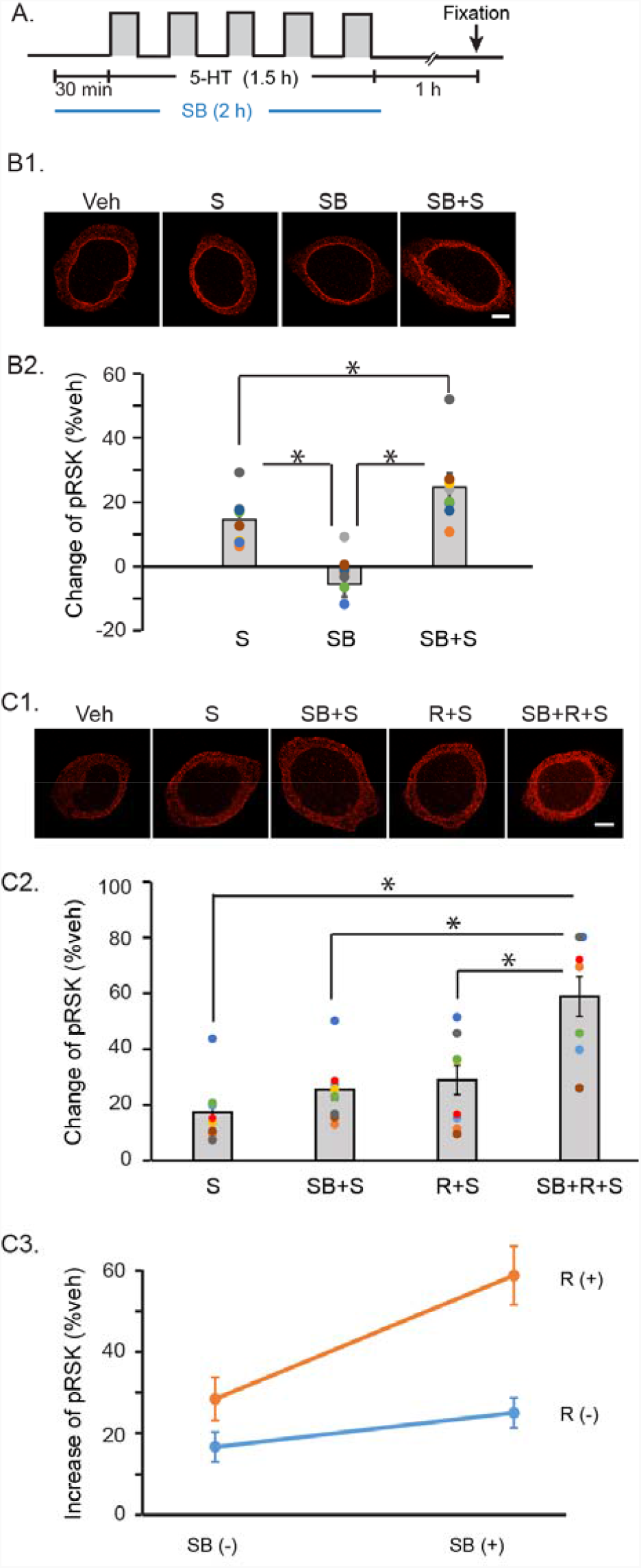
Effects of SB and rolipram on 5-HT-induced increase in pRSK. **A**, Protocol for standard 5 pulses of 5-HT treatment, in the present of p38 MAPK inhibitor SB 239063 (SB). **B**, Inhibition of p38 MAPK potentiates 5-HT-induced phosphorylation of RSK. **B1**, Representative confocal images of pRSK in SNs 1 h after 5-HT treatment, with SB. **B2**, Summary data. SB potentiated the 5-HT-induced increase in pRSK. **C**, Simultaneous activation of PKA and inhibition of p38 MAPK synergistically enhanced RSK activation. **C1**, Representative confocal images of pRSK in SNs 1 h after 5-HT treatment, with SB and/or rolipram. **C2**, Summary data. SB+R potentiated the 5-HT-induced increase in pRSK. C, Changes of pRSK in response to SB and rolipram (R). Two-way ANOVA revealed that R together with SB synergistically potentiated the 5-HT-induced increase in pRSK. Different colored dots in each group indicate individual experiments and the same color in different groups represent one experiment. Scale bar, 20 μm.

### 7. Activation of PKA and inhibition of p38 MAPK synergistically enhanced RSK activation

Modeling supported the hypothesis that co-application of a PKA activator and a p38 MAPK inhibitor would synergistically enhance RSK phosphorylation (Fig. 5C). We tested this hypothesis empirically. Rolipram and/or SB was applied to isolated SNs, and the effects on RSK phosphorylation were examined 1 h after 5-HT treatment. Immunoreactivities to pRSK in 5-HT, R+5-HT, SB+5-HT and R+SB+5-HT groups were normalized to vehicle control. Compared to Veh, 5-HT led to a 19± 4 % increase in pRSK. The increase in pRSK for R+5-HT and SB+5-HT was 25 ± 6% and 26 ± 5% of control, respectively. The increase of pRSK in the combined-drug group was 56 ± 8 % of control (Fig. 6C2). One-way RM ANOVA indicated significant overall differences among the groups (F_3.23_ = 22.13, p < 0.001). Post hoc comparisons (SNK) revealed the combined drugs induced a significant increase in pRSK, compared to the other 3 groups (SB+R+5-HT vs. 5-HT, q = 10.93, p < 0.001; SB+R+5-HT vs. SB+5-HT, q = 8.77, p < 0.001; SB+R+5-HT vs. R+5-HT, q = 7.89, p < 0.001), without significant effects from either SB or rolipram on 5-HT-induced pRSK (SB+ 5-HT vs. 5-HT, q = 2.25, p = 0.126; R+5-HT vs. 5-HT, q = 3.17, p = 0.085). Importantly, the interaction between rolipram and SB on pRSK was significant (Fig. 6C3) (Two-way RM ANOVA, F_1,7_ = 13.91, p = 0.007), indicating the drug combination of SB and rolipram synergistically increased 5-HT-induced phosphorylation of RSK.

## DISCUSSION

The reduction in LTF after RSK inhibition suggests *Aplysia* sensory-motor neuron co-cultures can be a useful cellular system for studying deficits in synaptic plasticity analogous to those associated with CLS.

### Synergism in rescue of deficits in LTF and in RSK activation

Drug combination therapies can take advantage of nonlinear properties of the signaling cascades mediating a biological response ^26,35,36^. This synergism can allow for lower drug doses which may reduce toxic drug side effects. Therefore, combination therapies are often superior to monotherapy^37-39^. It is plausible that combined-drug treatments similar to those we have examined could rescue deficient synaptic plasticity and memory such as that accompanying CLS, by simultaneously activating signaling pathways that converge onto a target (e.g. CREB1) necessary for long-term synaptic plasticity ^38,40-42^. Here we found that LTF impaired due to RSK inhibition by BID was unable to be restored by single drugs that individually activate either PKA or ERK pathways (Fig. 1). In addition, at these low doses, the drug combination had no adverse effects on basal synaptic strength (Fig. 2D), suggesting no side effects on synapses. However, the combined drugs at these doses produced a synergistic effect that rescued the disruption in synaptic plasticity in the CLS model (Figs. 2B, 3B). This rescue in BID-impaired LTF was accompanied by a synergistic increase in 5-HT-induced phosphorylated RSK (pRSK) in BID-treated SNs (Fig. 4C). This pRSK increase induced by dual drugs may explain, at least in part, the synergistic rescue of a downstream process, deficit LTF in the CLS model. It is also likely that the synergistic rescue of LTF also depends, in part, on actions of rolipram and NSC that do not involve RSK, such as enhanced phosphorylation of CREB1 by PKA, and enhanced phosphorylation of CREB2 by ERK, giving the evidence that dual drugs completely restored the impaired LTF in CLS model to normal enhanced LTF by dual drugs (Fig 2A and 2B).

### Computational modeling gives insight into determinants of synergism

The model of Zhang et al. (2021) was modified to explore conditions conducive to synergism by systematically simulating the effects of combined drugs on RSK and C/EBP. In this model (Fig. 5), the PKA-RSK feedforward path helps enhance RSK activity, whereas the ERK– p38 MAPK feedback loop suppresses the increase of pERK, reducing activity of RSK. The PKA and ERK / p38MAPK cascades interact to regulate the phosphorylation and activities of CREB1 and CREB2, which increase the synthesis and activation of C/EBP, leading to the increased pC/EBP essential for LTF.

Simulated effects of NSC or rolipram individually failed to induce strong synergism in both activation of RSK and normal LTF (Figs. 5B1-D), consistent with empirical results (Figs. 3A, 4B). The model predicted the negative feedback loop in which RSK activates p38 MAPK activation, which in turn inhibits MEK / ERK and thus RSK itself, may limit the extent of 5-HT induced RSK activation and LTF ^14^. Relieving this negative feedback would help exert synergistic effects. With RSK inhibited by BID, this negative feedback loop is weakened. Therefore, the range of potential RSK activation is less limited, which apparently facilitates synergistic interaction of NSC and rolipram (Fig. 5D).

The model also predicted that combining rolipram with a p38 MAPK inhibitor could be another useful drug combination, due to their synergistic activation of RSK and LTF (Figs. 5C-D). In contrast to BID that suppresses the activation of CREB1 (Fig. 5A, pathway 13), p38 MAPK inhibition suppresses activation of the transcription inhibitor CREB2 (Fig. 5A, pathway 15), which further enhances LTF. Potentiated activation of RSK by a p38 MAPK inhibitor was empirically verified (Fig. 6B) as was synergistic activation of RSK by a combination of this inhibitor with rolipram (Fig. 6C). Taken together, the above results demonstrate ways in which the complex interaction between PKA and MAPK pathways can affect the outcome of combined drugs. The intertwined positive feedforward cascade, which promotes synergism, and negative feedback loop, which suppresses synergism, together determined the dynamics of kinase activations and expression of transcription factors. Cellular models of disorders that affect synaptic plasticity and learning, such as CLS, may constitute a useful strategy to identify candidate drug combinations to treat aspects of these disorders, and combining biologically realistic computational models with empirical tests of predictions ^10,14,18,22,23^ can help explain synergism for drug combinations.

## Acknowledgments

The authors thank E. Kartikaningrum for preparing cultures. This study was supported by NIH grants NS019895 and NS102490.

## Author contributions statement

R. L., Y.Z., and J. B. contributed to the conception of the study. R. L. and Y. Z. designed and conducted the experiments and analyzed the results. P. S., L. C., and J. B. were involved in interpretation of the data and writing of the article. All authors reviewed the manuscript.

