## supplemental materials for "Defective synaptic plasticity in a model of Coffin-Lowry Syndrome is rescued by simultaneously targeting PKA and MAPK pathways"

**Equations and parameters of computational model**

Equations 1-40 are from the model of Zhang et al. (2021); equations 41-43 are modified from the model of Liu et al. (2013). Fourth-order Runge-Kutta integration was used for integration of all differential equations with a time step of 3 s. Further time step reduction did not lead to significant improvement in accuracy. The steady-state levels of variables were determined after at least two simulated days, prior to any manipulations. The model was programmed in XPPAUT (Ermentrout, 2002) (http://www.math.pitt.edu/~bard/xpp/xpp.html). Source codes will be submitted to the ModelDB database and to GitHub (https://github.com).

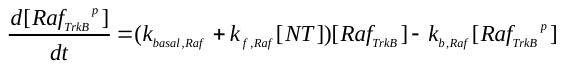
 (Eq. 1)

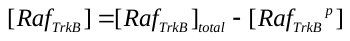
(Eq. 2)

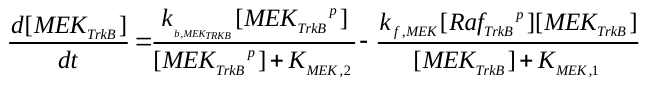
(Eq. 3)

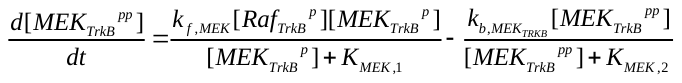
(Eq. 4)

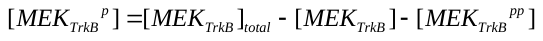
(Eq. 5)

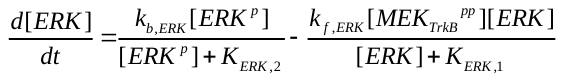
(Eq. 6)

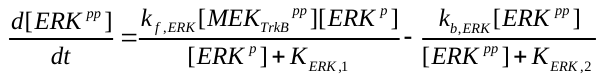
 (Eq. 7)

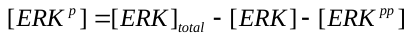
 (Eq. 8)

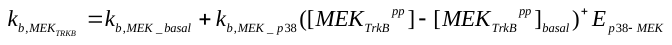
(Eq. 9)

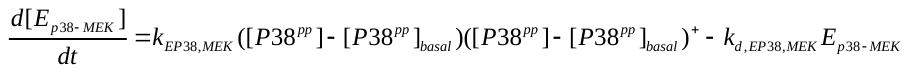
(Eq. 10)

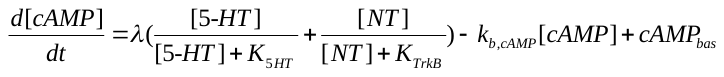
 (Eq. 11)

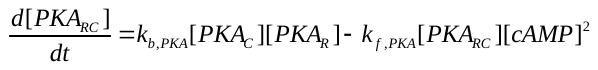
 (Eq. 12)

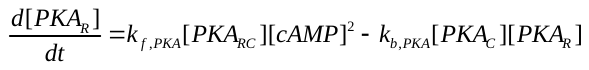
(Eq. 13)

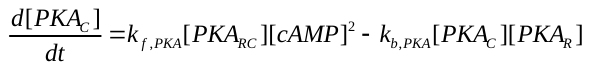
(Eq. 14)

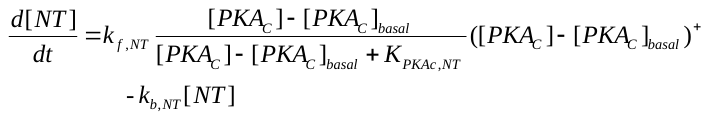
 (Eq. 15)

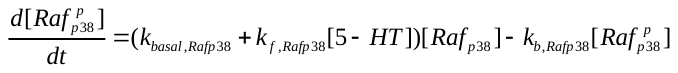
(Eq. 16)

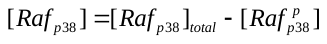
(Eq. 17)

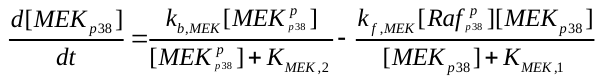
(Eq. 18)

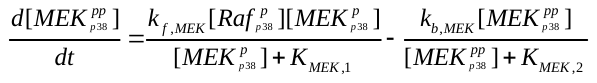
(Eq. 19)

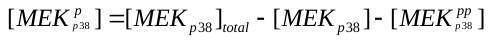
(Eq. 20)

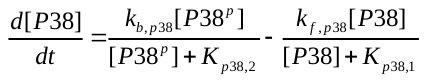
(Eq. 21)

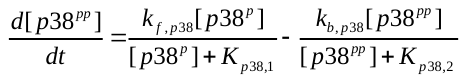
(Eq. 22)

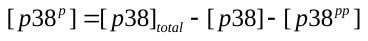
(Eq. 23)

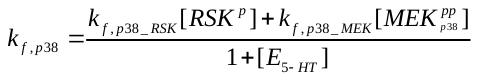
(Eq. 24)

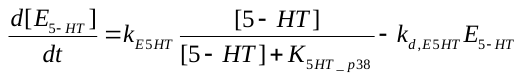
(Eq. 25)

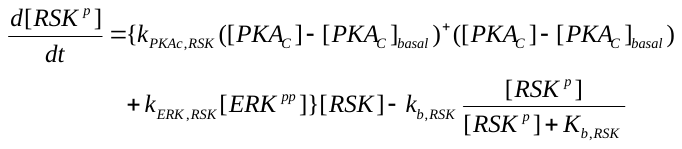
 (Eq. 26)

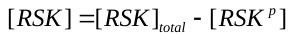
 (Eq. 27)

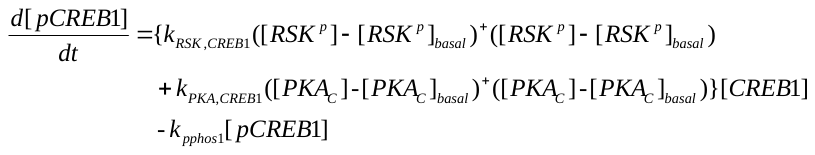
 (Eq. 28)
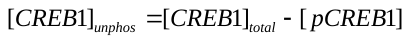
(Eq. 29)

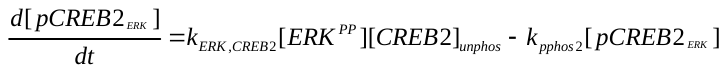
 (Eq. 30)

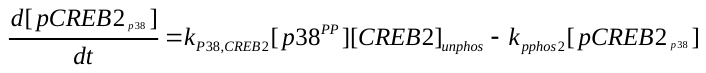
 (Eq. 31)

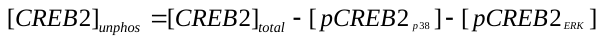
(Eq. 32)

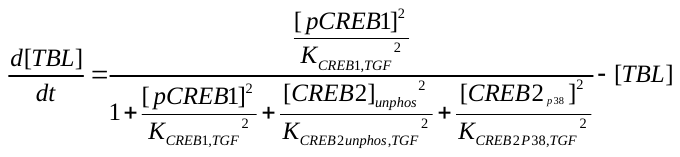
 (Eq. 33)

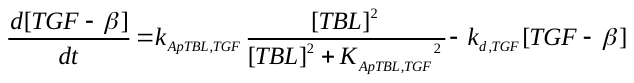
 (Eq. 34)

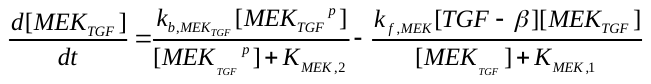
(Eq. 35)

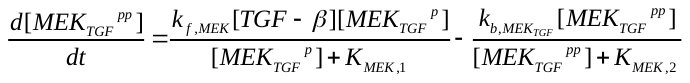
(Eq. 36)

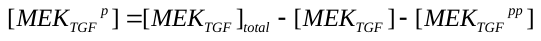
(Eq. 37)

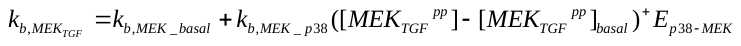
 (Eq. 38)

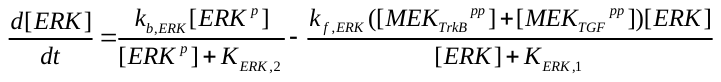
(Eq. 39)

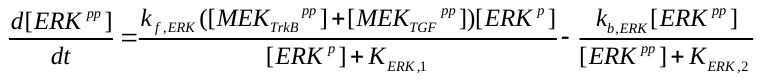
 (Eq. 40)

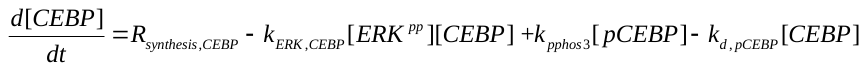
(Eq. 41)

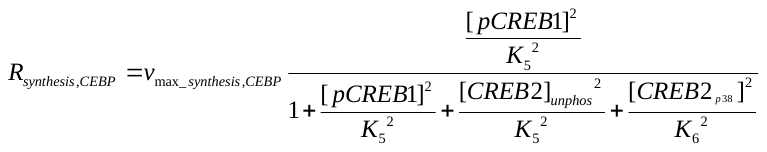
(Eq. 42)

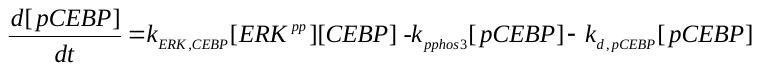
(Eq. 43)

Parameter values unchanged from the model of Zhang et al. (2021):

*k_f,MEK_* = 0.41 min^−1^, *k_b,MEK_basal_* = 0.04 μM/min, *k_b,MEK_p38_* = 0.04 min^−1^, *K_MEK,1_* = 0.20 μM, *K_MEK,2_* = 0.19 μM, *k_f,ERK_* = 0.41 min^−1^, *k_b,ERK_* = 0.12 μM/min, *K_ERK,1_* = 0.19 μM, *K_ERK,2_* = 0.21 μM, *[ERK]_tota_*_l_ = 0.5 μM, *k_basal,Rafp38_* = 0.036 min^−1^, *k_b,Rafp38_* = 0.1 min^−1^, *[Raf_p38_]_total_* = 0.5 μM, *[MEK _p38_]_total_* = 0.5 μM, *k_b,p38_* = 0.12 μM/min, *K_p38,1_* = 0.19 μM, *K_p38,2_* = 0.21 μM, *[p38]_tota_*_l_ = 0.5 μM, *K_5HT_p38_* = 50 μM, λ = 3.64 μM/min, *K_5HT_* = 85 μM, *k_b,cAMP_* = 1 min^−1^, *k_f,PKA_* = 20 μM^−2^min^−1^*,* *k_b,PKA_* = 12 μM^−1^min^−1^, *k_pphos1_* = 0.05 min^−1^, *[CREB1]_total_* = 0.05 μM, *k_ERK,CREB2_* = 3.5 μM^−1^min^−1^, *k_pphos2_* = 0.5 min^−1^, *[CREB2]_total_* = 0.05 μM, *k_f,Raf_* = 0.037 μM^−1^min^−1^, *k_basal,Raf_* = 0.00013 min^−1^, *k_b,Raf_* = 0.00038 min^−1^, *k_EP38,MEK_* = 0.33 μM^−1^ min^−1^, *k_d,EP38,MEK_* = 0.0013 min^−1^, *k_f,Rafp38_* = 0.0009 μM^−1^min^−1^, *k_E5HT_* = 0.8 min^−1^, *k_d,E5_HT_* = 1 min^−1^, *k_PKA,CREB1_* = 0.3 μM^−1^min^−1^, *[Raf_TrkB_]_total_* = 0.5 μM, *[MEK_TrkB_]_total_* = 0.5 μM, *[MEK_TGF_]_total_* = 0.083 μM, *[ERK^pp^]_basal_* = 0. 14 μM, *cAMP_bas_* = 0.7 μM, *[P38^pp^]_basal_* = 0.15 μM, *[MEK_TrkB_^pp^]_basal_* = 0.25 μM, *[MEK_TGF_^pp^]_basal_* = 0 μM, *k_f,p38_RSK_* = 0.99 min^−1^, *k_f,p38_MEK_* = 0.25 min^−1^, *K_TrkB_* = 12 μM, *k_f,NT_* = 0.3 μM/min, *[PKA_C_]_basal_* = 0.35μM, *K_PKAc,NT_* = 1 μM, *k_b,NT_* = 0.05 min^−1^, *K_b,RSK_* = 0.2 μM, *[RSK]_tota_*_l_ = 0.5 μM, *[RSK^p^]_basal_* = 0.042 μM, *k_PKAc,RSK_* = 0.26 μM^−1^min^−1^*, k_ERK,RSK_* = 0.82 μM^−1^min^−1^, *k_b,RSK_* = 0.31 μM^−1^ min^−1^, *k_RSK,CREB1_* = 1.25 μM^−1^min^−1^, *k_P38,CREB2_* = 3.5 μM^−1^min^−1^, *k_f,ApTBL_* = 1 μM/min, *k_b,ApTBL_* = 1 min^−1^, *K_CREB1,TGF_* = 0.003 μM, *K_CREB2unphos,TGF_* = 0.003 μM, *K_CREB2P38,TGF_* = 0.0003 μM, *K_ApTBL,TGF_* = 0.0078 μM.

Values of new parameters for equations 41-43:

*k_d,CEBP_* = 0.75 min^−1^, *k_ERK,CEBP_* = 0.25 μM^−1^min^−1^, *k_pphos3_* = 0.5 min^−1^, *v_max_synthesis,CEBP_* = 15.4 μM/min, *K_5_* = 0.003 μM, *K_6_* = 0.003 μM, *K_7_* = 0.0003 μM, *k_d,pCEBP_* = 0.0015 min^−1^

**Table S1.** Parameter values of the model.

**Supplemental Material & Methods**

Drug treatments

BI-D1870 (BID, Santa Cruz, #sc-397022A) is a small cell-permeant molecule that specifically inhibits mammalian RSK isoforms (RSK1, RSK2, RSK3 and RSK4) *in vitro* and in *vivo* by competitively inhibiting ATP at the N-terminal PKA/protein kinase G/protein kinase C domain of RSK. In this study, BID was used at concentration of 2 µM, which has been reported to block 5-HT-induced phosphorylation of CREB1, one of the RSK target proteins as well as LTF ^10^. To activate the ERK pathway, we used the DUSP6 inhibitor NSC 295642 (Sigma). At the concentration used, 0.01 μM, NSC specifically activates ERK but not p38 MAPK ^17^. To activate the PKA pathway, we used rolipram, an inhibitor of cAMP phosphodiesterase (PDE). At 0.2 μM, rolipram increases 5-HT-induced phosphorylation of CREB1 ^23^. However, 0.2 μM rolipram paired with 0.01 μM NSC interact to enhance basal synaptic strength without 5-HT stimulation. Moreover, NSC alone and rolipram alone potentiate 5-HT-induced weak LTF, and this weak LTF can be further enhanced by combining NSC and rolipram ^17^. To reduce the basal effects on synaptic strength from combined drugs, we decreased the concentration of rolipram to 0.1 μM. Rolipram at this concentration can still enhance 5-HT-induced phosphorylation of CREB1 (data not shown).
